# From Metabolism to Malignancy: Profiling Diabetes-Related Genes in Hepatocellular Carcinoma

**DOI:** 10.1101/2025.10.07.680510

**Authors:** Muhammad Sheraz, Danish Shakoor, Khizer Hayat, Hussam Tariq, Hasnain Khan, Muhammad Faraz Arshad Malik, Farhan Haq

## Abstract

Hepatocellular carcinoma (HCC) and diabetes mellitus both affect the liver, a key metabolic organ. This study uses bioinformatics to explore genetic links between Type 1 Diabetes (T1D) and HCC prognosis. Eleven HCC gene expression datasets from GEO and one TCGA-LIHC dataset were analyzed. Gene Set Enrichment Analysis (GSEA) identified up- and downregulated genes after data normalization in R. Four datasets (GSE64041, GSE78737, GSE107170, TCGA) revealed increased expression of T1D-related genes. Ten genes, including HLA-DOB and HLA-DPB1, were consistently upregulated. Statistical analysis (Kruskal-Wallis and Mann-Whitney U tests) showed these two genes were significantly associated with tumor grade and T-stage, with p-values ranging from 0.008 to 0.019. Co-expression analysis with 96 literature-curated T1D genes identified 23 related genes. Survival analysis using Kaplan-Meier curves highlighted five genes (IL7R, CD69, CCR5, RUNX3, PRF1), with CD69 showing strong associations with T-stage and disease-free survival. PRF1, RUNX3, and CCR5 were also linked to survival outcomes. Seven diabetes-related GEO datasets were used for validation. GSEA showed T1D gene enrichment in two datasets (GSE228267, GSE232310), with HLA-DOB significantly expressed in GSE228267 (p = 0.004). These findings suggest that HLA-DOB, HLA-DPB1, and three other T1D-related genes may serve as potential biomarkers for understanding the genetic connection between T1D and HCC. Though computational, this study lays the groundwork for future experimental validation.

## 1. Introduction

Liver cancer or Hepatocellular Carcinoma (HCC) is worldwide physical condition due to its high occurrence (Taniguchi H, 2023). Smoking, alcohol consumption, fatty liver and viral infections (Hepatitis B and C) (Kalantari, L., et al., 2023) are major risk factors for the high rate of HCC in developed countries. 80% of the total patients have a clinical history of HBV or HCV infection (Wang Y, et al., 2021). The genetic landscape of HCC is vast with *TP53* as a peak, regulating cell cycle and working as tumor suppressor protein where mutation leads to HCC (Yang Yang, et al., 2022). These genes are involved in regulation cells differentiation, proliferation and survival, hence indicate potential target for prognosis of HCC (Wang Y, et al., 2021). Some other genes that promote HCC are *CASP8, PIK3CA, APC, IGFR2*, and *AXIN1* (OMIM, 114550). The link between DM and cancer has been under investigation for 100 years. Diabetes Mellitus (DM) is another lethal metabolic disorder worldwide associated with abnormal insulin secretion or resistance to insulin pathways (Abubakar et al., 2021). Statistics of *International Diabetes mellitus Association* (IDF) show that, in 2021, 537 million adults (20-79 years) are living with DM which accounts for 1 in 10 people. This quantity is forecasted to rise to 643 million by 2030 and 783 million by 2045. Type II DM(T2DM), which is characterized by the body’s failure to use insulin, justify more than 95% of the total cases. In contrast, Type I DM(T1DM), which is characterized by less or no insulin production, accounts for less than 5% of the total cases (WHO, 2023). The genetic inheritance of T2DM is Autosomal dominant and some genes that are involved are *IRS1, PPARG, IL6, TCF7L2, AKT2, HMGA1* and *PAX4*(OMIM, 125853). T1DM inheritance is more complex and is not like other Mendelian diseases (Redondo et al., 2022). 30-35% of the T1DM is caused by mutations in *HLA* Class I (*HLA-A, HLA-B, HLA-C*) and *HLA* Class II *(HLA-DR, HLA-DQ, HLA-DP*) genes (Bell, G.I et al., 2022). A few replicated genes loci of insulin gene *INS* and some immune related genes; *CTLA4, PTPN22* and *IL2RA* have a role in T1DM (Bottini, N et al., 2004).

DM and HCC are complexly related to each other through common risk factors like obesity, hyperglycemia and insulin injections. Cancer risk appears to be higher for both T1DM and T2DM patients, with T1DM accounting for around 8-18% of cancer patients (Zhu Bing and Qu Shen, 2022). Co-occurrence of HCC with T2DM is more frequent yet the underlying mechanism remains unclear (Hongpeng Jiang, et al., 2022). Insulin resistance and insulin receptor activation pathways are involved in the progression of HCC (Plaz Torres, et al., 2022). Several findings have demonstrated many genes that are associated with both DM and HCC including *CCNA2, MAD2L1, BUB1, RACGAP1, NCAPG* and *ASPM* (Shi, Z., et al., 2020). In this analysis we are directing to study diabetes and HCC through genomic investigation of Datasets from Gene Expression Omnibus (GEO) using Gene Set Enrichment Analysis (GSEA).

## 2. Methodology

### 2.1 Data retrieval

As a base for careful genomic study, appropriate datasets were analytically acquired from the Gene Expression Omnibus (GEO) database. The datasets considered for investigation include GSE107170 (Diaz et al., 2018), that includes 307 samples (134 tumor, 144 normal, and 29 cirrhosis); GSE102079 (Chiyonobu et al., 2018; Hatano et al., 2023), having 257 samples (13 normal and 242 tumor); GSE78737 (Sekhar et al., 2018), which include 107 samples (103 transplant and 4 hepatocellular carcinoma); and GSE69715 (Sekhar et al., 2018), having 103 samples (66 normal and 37 tumor), GSE41804 (Hodo et al., 2013), including 40 samples (20 normal and 20 tumor); GSE25097 (Tung et al., 2011; Lamb et al., 2011; Sung et al., 2012; Wong et al., 2016; Srivastava et al., 2012; Ivanovska et al., 2011), having 557 samples (289 normal and 268 tumor); and GSE15765 (Woo et al., 2010; Woo et al., 2008), which include 90 samples (77 tumor and 13 cholangiocarcinoma). Likewise, an additional dataset from TCGA PanCancer was retrieved via cBioPortal (Cerami et al., 2012; de Bruijn et al., 2023; Gao et al., 2013), on Liver Hepatocellular Carcinoma, ensuring a diverse dataset selection for further studies.

### 2.2 Data Normalization and Differential Expression

To reduce any bias or variations and maintain the integrity of our data, raw data collected from GEO is preprocessed prior to any investigation. We used TPM (Transcript Per Million) for RNA-seq data normalization or RPKM (Reads Per Kilobase per Million) are used while microarray data is normalized through RMA (Robust Multi-Array Average). We used R software to normalize the datasets. Differential expressions are estimated using limma packages. Genes that show no or low expression are filtered out leaving us with only significant genes.

### 2.3 Gene Set Enrichment Analysis (GSEA) setup

After normalization, datasets are processed through GSEA software. It focuses on entire gene sets rather than individual genes. For each dataset of diabetes and HCC, gene expression profiles are ranked according to statistics used in GSEA (fold change, p value or enrichment score) between diseased state and control group. The ranked genes are compared to predefined gene sets such as KEGG or Reactome, that are functionally related to insulin signaling, inflammation, cell cycle regulation and lipid metabolism.

### 2.4 Statistical analysis

R or SPSS is used to conduct all statistical analyses, including clinical analysis and survival analysis. To generate Kaplan-Meier survival plots of significantly enriched or expressed genes R version 4.2.2 is used. A Mann Whitney U-test and Kruskal Wallis tests were applied using IBM SPSS 21 software (IBM Inc., Armonk, NY, USA). p-values < 0.05 were considered statistically significant. GraphPad Prism 5 was used to draw out the graphical representation of the data.

## 3. Results

GSE107170 and TCGA showed significant enrichment of genes linked with Type 1 Diabetes (T1D), with highly significant p-values and enrichment scores (ES) among multiple comparisons. In GSE107170, T1D genes were substantially more abundant in tumor samples than in normal tissues (P = 0.008, ES = 0.76). Similarly, GSE78737 was significantly overrepresented in transplant patients compared to normal samples (P = 0.015, ES = 0.75). GSE64041 revealed a substantial enrichment in HBV-associated hepatocellular carcinoma compared to HBV-infected non-cancerous tissues (P = 0.034, ES = 0.68). Furthermore, the TCGA dataset revealed a substantial enrichment in advanced-stage cancers (T3,4) versus early-stage tumors (T1,2) (P = 0.03, ES = −0.69).

The evaluation of T1D-associated genes within the selected datasets found that GSE64041 had 26 related genes, GSE78737 had 16, GSE107170 had 14, and the TCGA dataset had 26 genes with core enrichment designated as YES. All datasets showed core enrichment toward the very top of the ranking gene lists. However, in the TCGA dataset, there was no early core enrichment, but 26 genes near the end of the sorted list showed core enrichment, suggesting a negative enrichment pattern. This shows that these genes may be downregulated in tumor samples or linked to unique regulatory mechanisms in hepatocellular carcinoma progression. Only ten of the discovered genes were consistently present in all four datasets: CD86, IL1B, and HLA genes (DMA, DQB1, DMB, DRA, DPB1, DQA1, DOB, and DRB1).

**Figure. 1:**
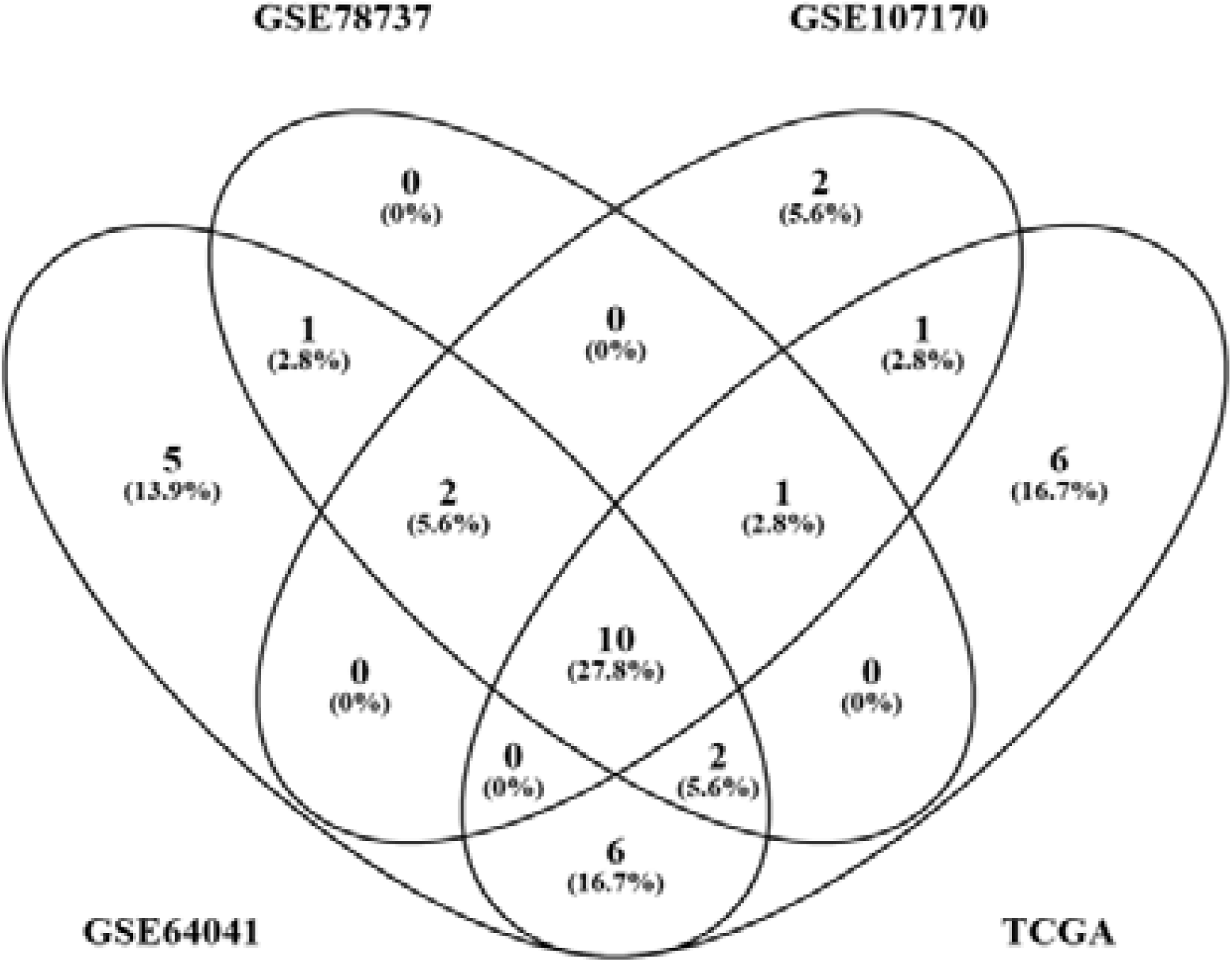

Correlation of these 10 genes was calculated using R software and heatmaps were created.

**Figure. 2:**
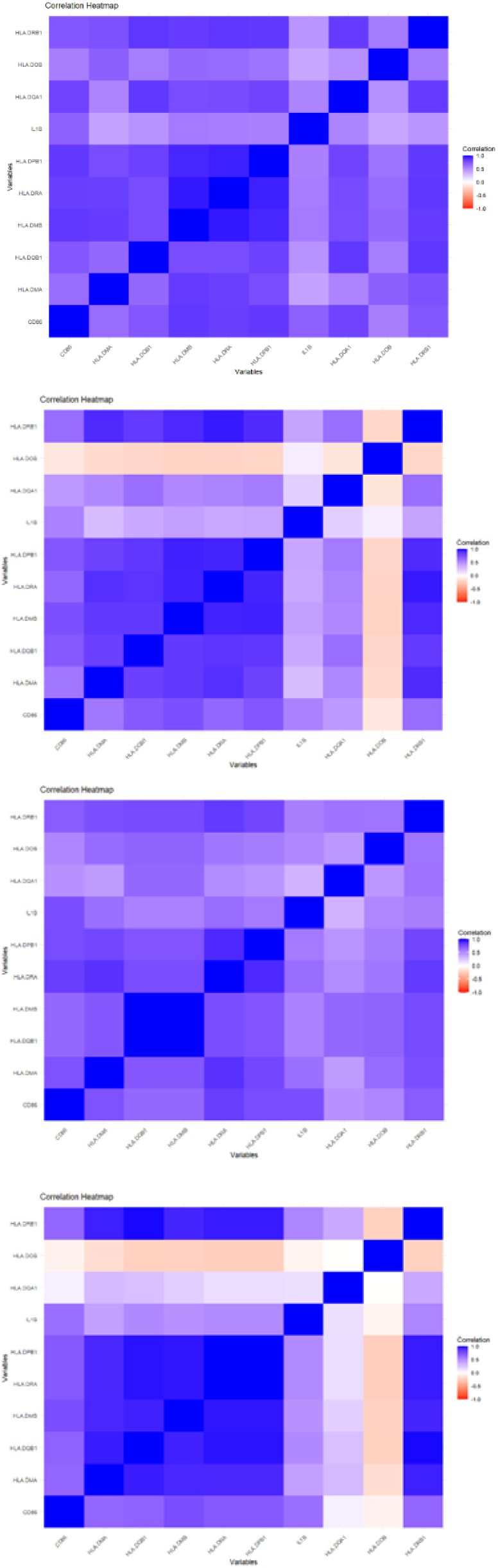

Mann Whitney and Kruskal Wallis Tests were applied to these 10 genes using different clinical parameters like Tumor stages, metastasis occurrence, lymph node stage and T stages from the dataset of TCGA. Out of these 10 genes 2 of the genes viz HLA-DOB and HLA-DPB1. HLA-DPB1 is significant in Early/Advance tumor grade with p=0.01 and Early/Advance T Stage with p=0.013 and HLA-DOB is significant in Early/Advance tumor grade and Early/Advance T Stage with p values of 0.008 and 0.019 respectively.

**Figure. 3:**
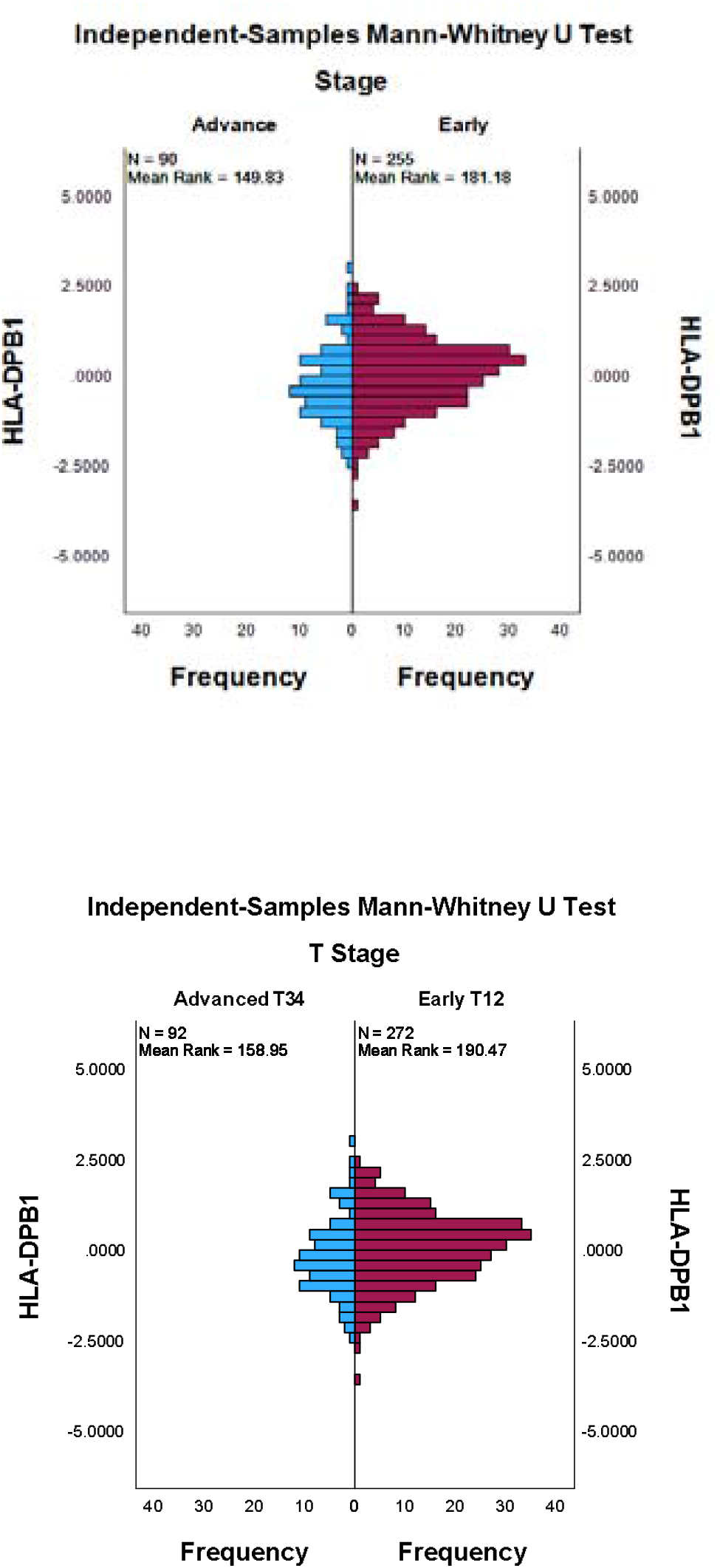

**Figure. 4:**
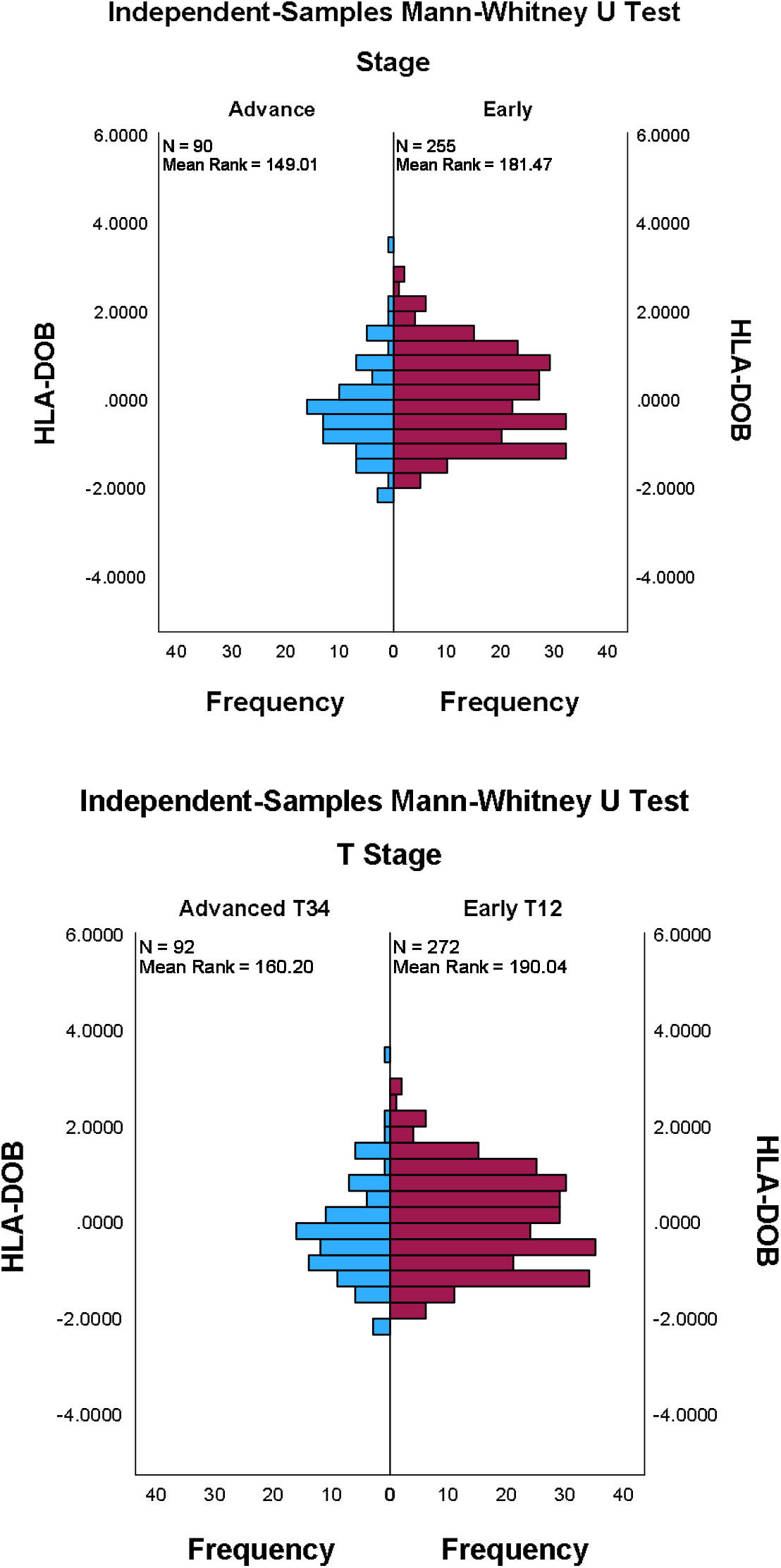

Redondo et al.’s (2023) analysis revealed 98 genes related with diabetes. The Z-scores of these genes were taken from The Cancer Genome Atlas (TCGA) dataset, together with the Z-scores of HLA-DOB and HLA-DPB1, to determine the relationship amongst these two hepatocellular carcinoma (HCC)-associated genes and diabetes-related genes. To analyze their relationships, Pearson correlation analysis was implemented with IBM SPSS. The findings revealed a strong link between HLA-DOB and HLA-DPB1 and 22 type 1 diabetes (T1D)-related genes identified in the literature. These genes are IRF1, FASLG, IKZF1, CCR5, RASGRP1, RUNX3, PTPN22, CCDC88B, PRF1, CD69, PTPRC, IL2RA, RGS10, IRF4, GATA3, IL10, GPR183, CAMK4, ITGB7, FLI1, IL7R, and IKZF3.

**Figure. 5:**
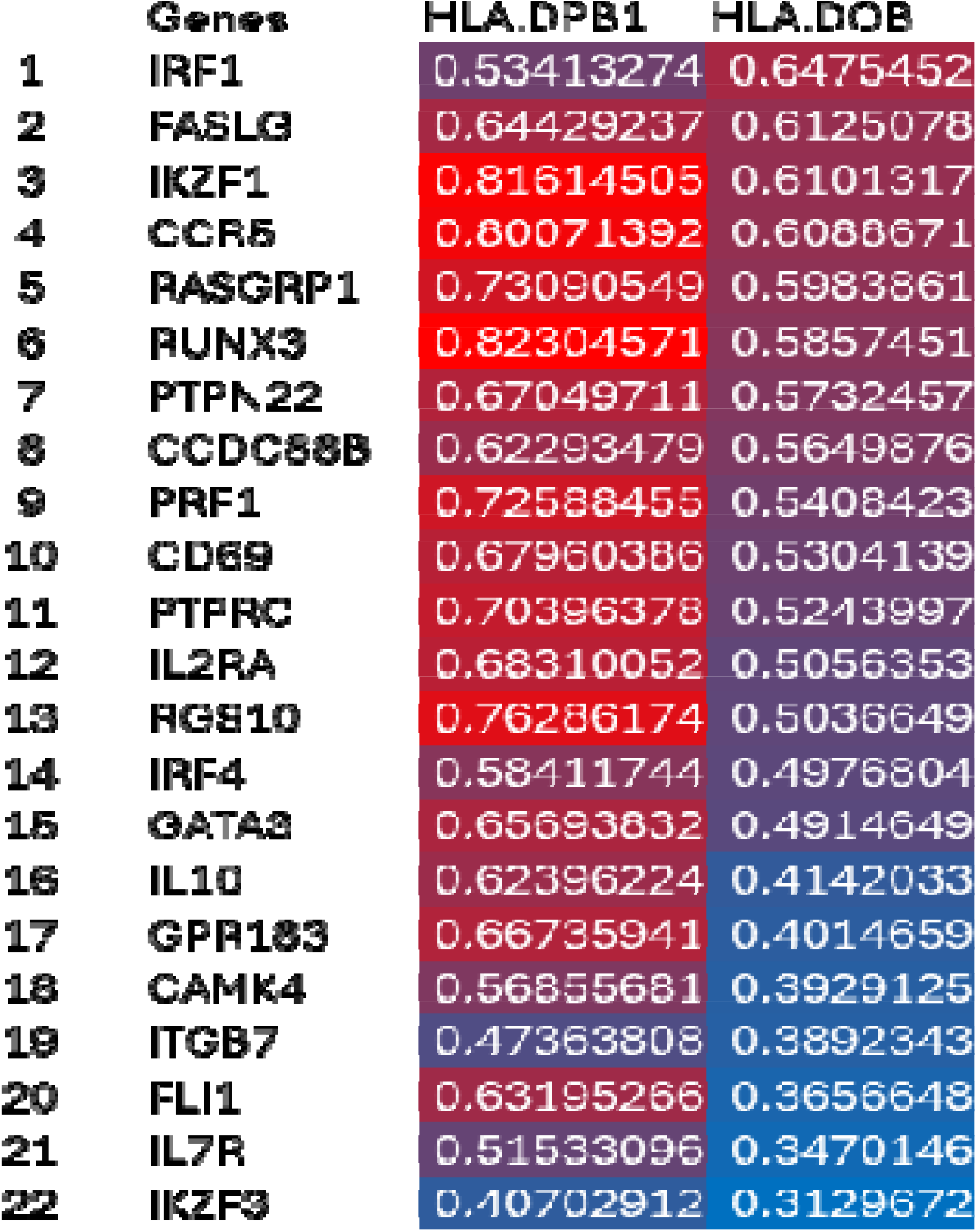

The Z-scores of these 22 genes were binarized for disease-specific survival analysis with the Kaplan-Meier survival test. Genes with Z-scores equal to or above the median were given a value of one, while those with Z-scores less than the median were assigned a value of zero. According to the Kaplan-Meier survival study, only five genes, RUNX3, PRF1, IL7R, CD69, and CCR5, were significantly related to disease-specific survival with p values of 0.006, 0.032, 0.014, 0.049, and 0.039 respectively

**Figure. 6:**
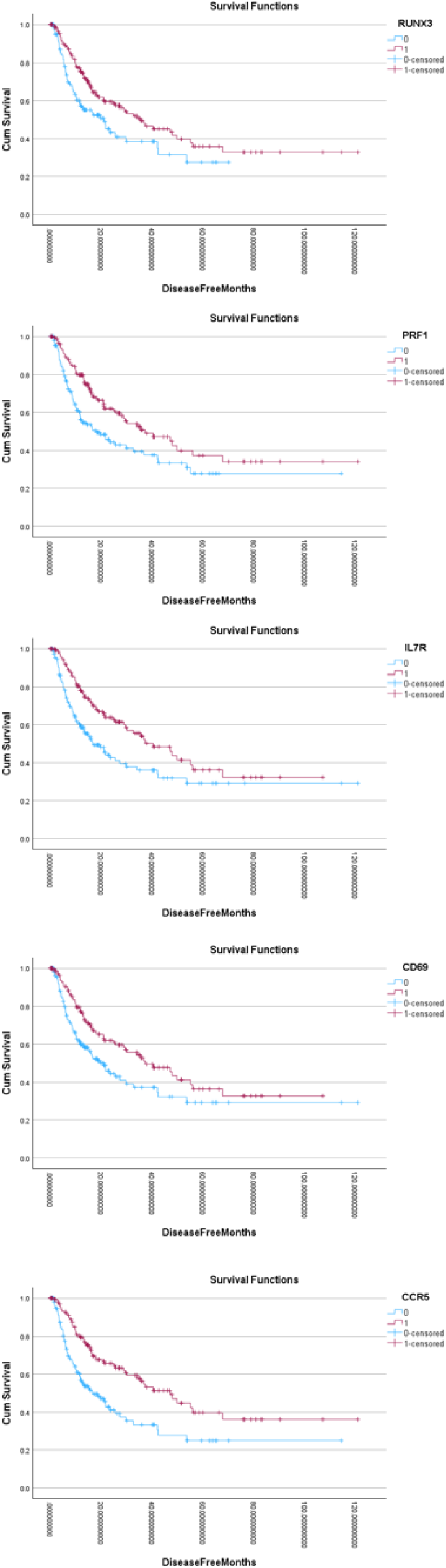

Clinical analysis of these five important genes was performed using the Kaplan-Meier survival test, with the same classification criteria, assigning a value of 1 to genes with Z-scores equal to or above the median and 0 to those below the median. The results showed that all five genes had substantial statistical significance: RUNX3 (P = 0.011), PRF1 (P = 0.003), IL7R (P = 0.002), CD69 (P = 0.009), and CCR5 (P = 0.001).

**Figure. 7:**
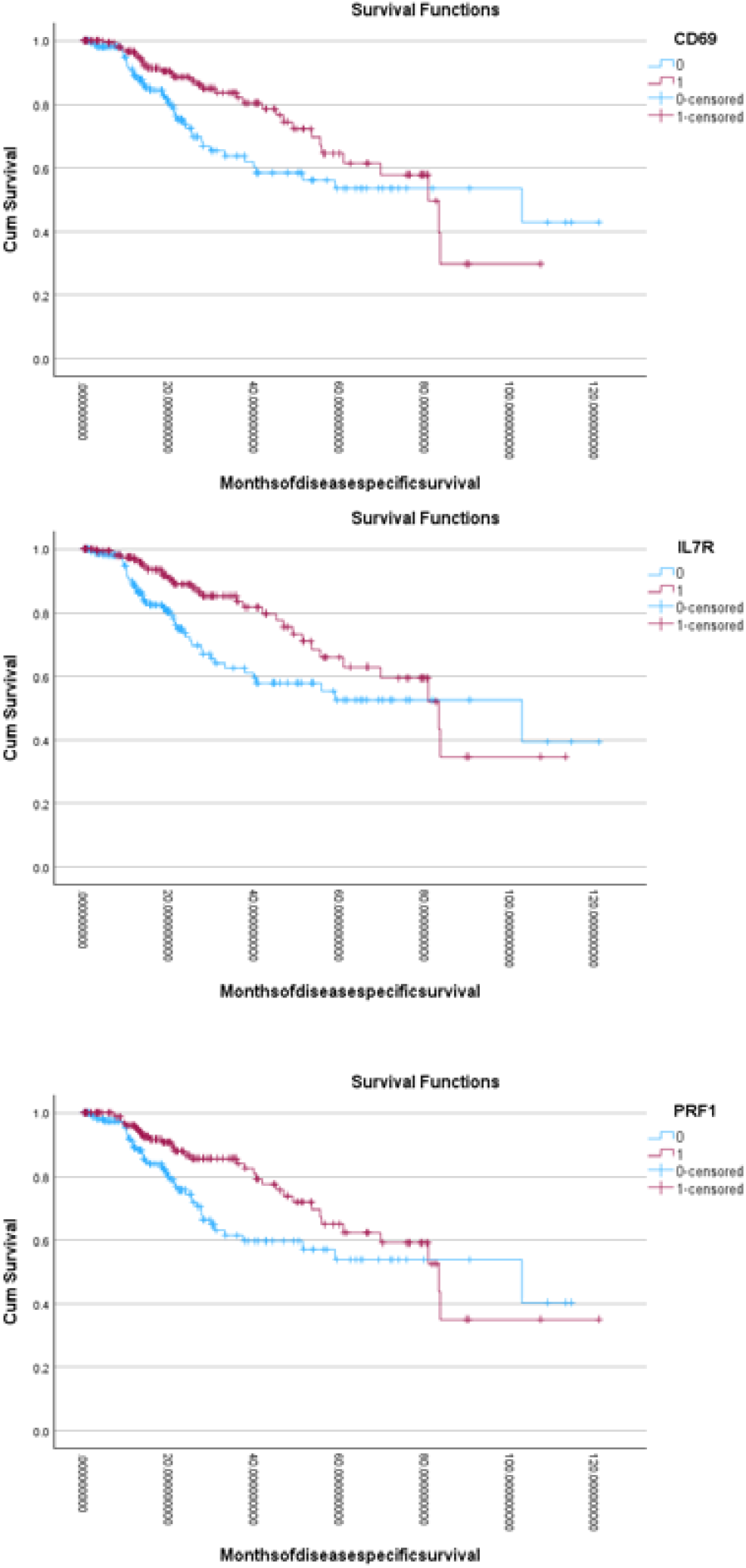

**Figure. 8:**
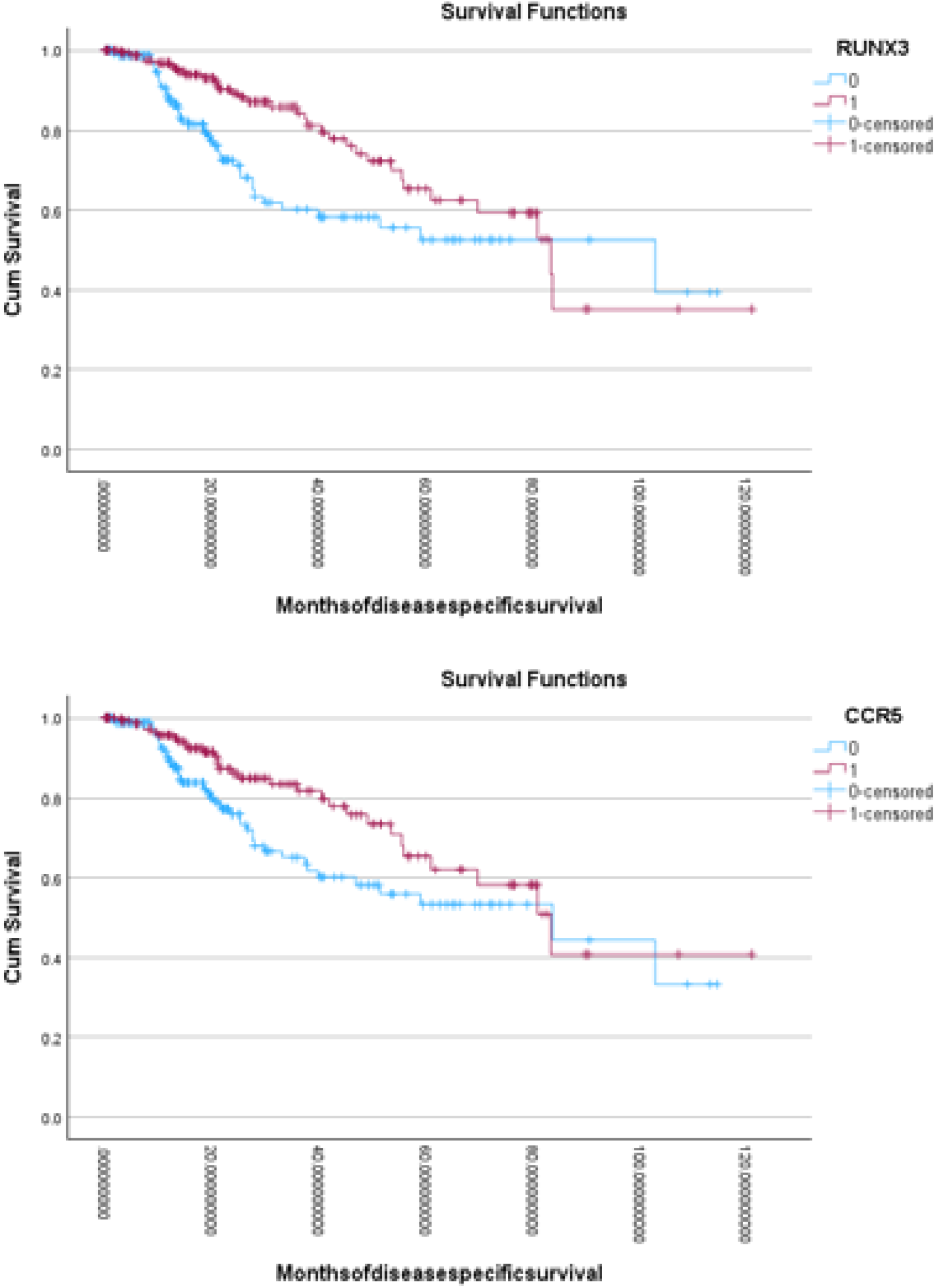

## 4. Discussion

This work establishes a significant molecular relationship between Type 1 Diabetes (T1D) and Hepatocellular Carcinoma (HCC) by finding important genes and pathways shared by both disorders. We found considerable enrichment of T1D-associated genes in many datasets (GSE107170, TCGA, GSE78737, and GSE64041), with significant p-values and high enrichment scores. Notably, T1D-related genes were more prevalent in tumor samples than in normal tissues, particularly in advanced-stage HCC (T3,4), indicating a potential role in cancer progression. However, a negative enrichment pattern in the TCGA data suggests that these genes are downregulated in tumor samples or are involved in unique regulatory pathways that may influence HCC formation.

Further analysis revealed ten consistently present genes, including HLA-DOB and HLA-DPB1, both of which had significant relationships with tumor growth. These genes were significantly related to identifying early versus advanced tumor grades and T-stages, indicating their potential as HCC biomarkers. Furthermore, Pearson correlation analysis found a strong link between HLA-DOB and HLA-DPB1 and 22 diabetes-associated genes, confirming the theory that metabolic and immunological dysfunctions in diabetes contribute to HCC development.

A Kaplan-Meier survival study of these 22 genes identified five genes—RUNX3, PRF1, IL7R, CD69, and CCR5—that were significantly related with disease-specific survival. These results are uniquely relevant when considering the pathways linked with these genes, which are concerned in both the development of Type 1 Diabetes and Hepatocellular Carcinoma. RUNX3 is affected in cell-cycle control via CDKN1A transcription, as well as immunological responses and cell migration. It also has a significant role in NOTCH, Wnt, and TGF-beta, which have all been linked to tumor growth and immune modulation in both T1D and HCC. Th1 and Th2 cell development are also regulated by RUNX3, which shows its role in immune response in both T1D and HCC.

PRF1, another survival-associated gene, that plays a role in natural killer (NK) cell-mediated cytotoxicity, critical for immune surveillance and tumor cell reduction. Pathways related to PRF1 are linked to apoptosis, autoimmune disorders, and the immunological response in T1D, and they interconnect with pathways like FoxO signaling, PI3K-Akt signaling, and JAK-STAT signaling, that are all associated with tumor progression.

CCR5 is another important gene that is involved in immune cell migration and has been linked to a diversity of inflammatory disorders. HIV, toxoplasmosis, and viral carcinogenesis are all examples of CCR5-associated pathways that can promote cancer development. CCR5’s involvement in the chemokine signaling system and viral carcinogenesis, proposes a role in the inflammatory developments that link diabetes and HCC. The cytokine-cytokine receptor interaction pathway is very important during T1D and HCC, that supports its function as a potential biomarker.

These five genes were validated using clinical data, resulting highly significant p-values, supporting their great predictive relevance. Their involvement in immunological and inflammatory pathways emphasizes the crucial role of immune system disorders and metabolic abnormalities in the progression of HCC in diabetics.

Gene Set Enrichment Analysis (GSEA) was executed on seven datasets from the Gene Expression Omnibus (GEO) to specify additional validation. The GSE228267 dataset (De Jesus et al., 2024) resulted a significant enrichment of the T1D gene set with a p-value of P = 0.004, and this dataset included HLA-DOB. Another dataset, GSE232310, had a non-significant p-value of P = 0.142, however HLA-DOB was still enriched, demonstrating its importance.

Diabetes mellitus (DM) and HCC are intricately linked by common risk factors such as obesity, hyperglycemia, and insulin therapy. Shi et al. (2020) reported 256 genes in both DM and HCC, in which 155 were elevated and 101 were downregulated. Severe obesity, a typical risk factor of Type 2 Diabetes, has been associated to elevated death rates in HCC (Calle et al., 2003), which is caused by processes such as cell proliferation, hormone imbalance, and inflammation. Hyperglycemia is another important factor in carcinogenesis, as reported by Zhu Bing and Qu Shen (2023), who discovered a significant rise in tumor prevalence, including HCC, in hyperglycemic patients.

Devic (2016) defines the Warburg effect as cancer cells relying on glycolysis rather than mitochondrial phosphorylation for energy, resulting in increased glucose uptake and cell proliferation. Furthermore, insulin therapy, particularly insulin analogs like lispro, aspart, and glargine, have been linked to an increased risk of cancer due to their carcinogenic effects and higher IGF-1 levels (Hemkens et al., 2009). Metformin, an anti-diabetic medication, has shown hepatoprotective effects and a lower risk of cancer. Zhang et al. (2013) and Singh et al. (2013) discovered that metformin use dramatically reduced the incidence of HCC, perhaps via lowering hyperinsulinemia and inhibiting the mTOR pathway.

According to studies, diabetic livers are 2.5 times more likely to develop HCC than non-diabetic livers (Bhrigu Kumar and Pramod C, 2021). Insulin receptors are overexpressed in HCC, and insulin’s interaction with hepatocytes activates the Akt pathway, encouraging tumor growth. Teng P et al. (2023) noted that DM contributes to hepatocarcinogenesis by activating inflammatory pathways that result in the formation of reactive oxygen species, proinflammatory cytokines, and tumor growth.

## 5. Conclusion

In summary, this study sheds light on the molecular pathways that diabetes and HCC overlap, emphasizing the vital roles that immunological and metabolic dysregulation play in the development of cancer. Finding HLA-DOB, HLA-DPB1, and the five survival-associated genes opens exciting possibilities for finding biomarkers and creating focused treatment plans. These results highlight the potential of customized therapy to improve treatment outcomes and disease prognosis among individuals having both diabetes and HCC.

## Notes

### Competing Interest Statement

The authors have declared no competing interest.

